# Synthesis and characterization of poly(propylene imine) dendrimers, as nanocarriers of Benznidazole: an in vitro controlled release assay

**DOI:** 10.1101/2022.07.20.500757

**Authors:** Jenny Ordoñez-Benavides, Henry Andrade-Caicedo

**Affiliations:** Grupo de Investigación en Dinámica Cardiovascular, Centro de Bioingeniería, Escuela Ciencias de la Salud, Universidad Pontificia Bolivariana, Medellín, Antioquía, Colombia; Facultad de Ciencias Exactas y Aplicadas, Instituto Tecnológico Metropolitano ITM, Medellín, Antioquía, Colombia; Grupo de Investigaciones en Bioingeniería y Microelectrónica, Centro de Bioingeniería, Escuela de Ingenierías, Universidad Pontificia Bolivariana, Medellín, Antioquía, Colombia

**Keywords:** Poly(propylene imine), Nanocarrier, Benznidazole, Chagas disease, Synthesis, Characterization, *Trypanosoma cruzi*, Loading efficiency, Drug encapsulation

## Abstract

**Background:** American trypanosomiasis, or Chagas disease, is the result of an infection caused by the *Trypanosoma cruzi parasite*. The disease is endemic in Latin America, where the main clinical manifestation and cause of death of Chagas patients is cardiomyopathy. The current approved treatment for this disease is based on the use of the nitroheterocyclic compound, Benznidazole. The drug is administered in high doses and for prolonged periods, which causes serious adverse effects, eventually leading to treatment discontinuation. In addition, it has only shown efficacy in the acute phase of the disease. Benznidazole has low solubility, low permeability, low bioavailability and high toxicity in the body. These physicochemical characteristics can be improved by using dendritic structures that serve as nanocarriers.

**Methods:** In this research, poly(propylene imine) PPI dendrimers in generations 4.0 G and 5.0 G were synthesized and characterized. We performed the synthesis by divergent approach. We encapsulated Benznidazole using the equilibrium dialysis method, and we evaluated the loading efficiency and the concentration of the released drug by high-performance liquid chromatography (HPLC).

**Results:** Preliminary results showed a drug loading efficiency on the dendrimer of 78% and an entrapment percentage of 99.6%. The release kinetics showed a controlled and sustained release over time compared to dendrimer-free Benznidazole.

**Conclusions:** The PPI 5.0 G - Benznidazole dendrimer system could be considered as an alternative to be evaluated in vitro and in vivo, as an alternative to conventional treatment of Chagas disease. The next stage of the experimental work consists of standardizing an infection model of H9C2 cardiomyocytes with Colombian strains of *Trypanosoma cruzi*, in order to evaluate the effect of the encapsulated drug on nanocarriers.

## Introduction

Chagas disease is a serious and important public health problem in Latin America, both in terms of health, socioeconomic impact and geographical distribution (1)(2). It is estimated that eight million people are infected worldwide, mainly in Latin America, where the disease is endemic in 21 countries (3)(4). Each year, 41,200 new cases of infected persons are reported, approximately 12,000 deaths per year due to cardiac complications, and 586 000 young adults of productive age are disabled. Currently, millions of chronically infected people are at risk of developing cardiovascular and/or digestive pathologies, making Chagas disease one of the main causes of morbidity and premature death due to cardiac complications in Latin America (5)(6). In addition, migratory phenomena have caused the expansion of the disease to non-endemic countries such as the United States, Canada and several European countries, Japan and Australia (5). The therapy currently available for chagasic patients at any stage is limited to the use of the FDA-approved orally administered nitroheterocyclic drug Benznidazole (BZN). The drug has been shown effective in the acute phase of the disease, where the parasitological cure is 80%. However, in the indeterminate and chronic phases, only about 5 to 10% parasitological cure has been demonstrated (2).

Benznidazole is hydrophobic in nature, therefore, it has low permeability, low solubility, low bioavailability and high toxicity in the body and can be rapidly metabolized by the liver, leaving very little active ingredient in the specifically injured tissue. The Biopharmaceutical Classification System has assigned a class IV categorization, which is due to the low solubility of the drug in water and its low permeability (7). These properties lead to the drug impacting uninjured tissues causing side effects, making the treatment ineffective and unsafe (8). Regarding the therapeutic management of chronic Chagas cardiomyopathy (CCC), in addition to antiparasitic treatment, it has been managed according to the conventional institutionalized treatment for similar cardiomyopathies caused by other etiologies (9). The antiparasitic treatment could be improved if some of the physicochemical properties of the trypanocidal drug, such as solubility, bioavailability and bioaccumulation, are modified and/or improved. Currently, several researches have been developed to improve drug solubility. These investigations are aimed at using techniques such as solid dispersion (10,11), nanosuspension methods (12)(13), particle size reduction (14)(15), cryogenic methods (16), micellar solubilization techniques (17)(16), processes with supercritical fluids (18), and the use of dendrimers (19) (20). Among these methods, the use of dendrimers has gained much attention due to the structure, surface functional groups and presence of electrostatic and covalent bonds between the drug and the dendrimer (20). The dendrimer-drug complex improves bioavailability, release time and drug delivery in the injured tissue (21). Dendrimers are spherical, monodisperse nanostructures with a symmetrical structure, having a central core surrounded by branches composed of repeating monomeric units and terminal functional groups. These structural characteristics allow drugs to be encapsulated in the core or conjugated to the functional groups (22). Dendrimers offer the advantage of encapsulating the drug, and additionally, they allow the conjugation of molecular targets and biocompatible coating molecules in their terminal functional groups, forming polymeric nanoplatforms, making therapy increasingly effective and selective (23).

Regarding the use of dendrimers in parasitic treatment, few results have been published. Giarolla et al., demonstrated the possibility of using dendrimers as drug transporters. They constructed dendrimers from myoinositol, D-mannose and malic acid. The dendrimer was used to encapsulate hydroxymethylnitrofurazone as a bioactive agent. Fernandez et al., evaluated the use of a first-generation dendrimer as a candidate for delivery of the anti-*T. cruzi* compound, (2′-(benzo [1,2-c] 1,2,5-oxadiazole-5(6)-yl (N-1-oxide) methylidene]-1-methoxy me- thane hydrazide), finding that dendrimers could be suitable for transporting antichagasic drugs. The research of Garolla and Fernández is reported in the work of Juárez-Chavéz et al. (24).

Among the dendrimers that can be used for drug release are poly(propylene imine) (PPI) dendrimers. PPI dendrimers are synthesized by divergent approach, by double Michael addition reaction of acrylonitrile to ethylenediamine nuclei, followed by a reduction of CN groups to primary amino groups by catalyst-assisted hydrogenation (25).

In this research work, we synthesized PPI dendrimers in generations 4.0 G and 5.0 G by divergent approach, using ethylenediamine as core and acrylonitrile as branches. The synthesized dendrimers were characterized both physicochemical and morphologically by infrared spectroscopy (FTIR), proton nuclear magnetic resonance (HMNR), transmission electron microscopy (TEM) and nanoparticle tracking analysis (NTA). We used the synthesized dendrimers to encapsulate BNZ. Finally, we evaluated the drug loading into the dendrimer and the release kinetics of the encapsulated drug in vitro by HPLC. The concentrations were determined from the calibration curve of the pure drug.

## 2. Materials and methods

### 2.1 Materials

Benznidazole, (N-benzyl-2-nitro-1-imidazolacetamide) was donated by the American Trypanosomiasis Laboratory of the Institute of Biomedical Research of the National Autonomous University of Mexico (UNAM). Ethylenediamine (C_2_H_8_N_2,_ 99% purity from PANREAC), acrylonitrile (C_3_H_3_N, 99% purity from Sigma – Aldrich), gaseous hydrogen (H_2_ UAP grade 5 from CRYOGAS), Raney Níquel (Merck), methanol (CH_3_OH 99% purity from Sigma – Aldrich), phosphate-buffered saline solution PBS 0,01M, pH 7,4 (SIGMA, USA), cellulose membrane 12-14 kDa (spectral/by molecular porous membrane tubing, USA), acetonitrile HPLC grade (Merk). The chemical reagents used in the synthesis were analytical grade and HPLC grade.

### 2.2 HPLC analytical method for Benznidazole quantification

We quantified BZN according to the method reported by Da Silva RM (26) with some modifications. We used the HPLC (SHIMADZU LC-2010), available at the biology laboratory of Universidad Pontificia Bolivariana and an LC 18 column, 250*4.6 mm, 5 µm, with an operating temperature of 25°C. The mobile phase consisted of a mixture of acetonitrile HPLC grade and ultrapure water (5:95), which was pumped with an isocratic flow rate of 0.7 ml/min. BZN was detected by monitoring the absorbance of the eluent column at 324 nm in an Ultraviolet-Visible detector. The concentrations of the analyte were determined from a calibration curve of the pure analyte.

### 2.3 Synthesis and characterization of poly (propylene imine) dendrimers (PPI)

We synthesized PPI dendrimers in generations 4.0 G and 5.0 G by double Michael addition reaction of acrylonitrile to ethylenediamine cores, followed by heterogeneous hydrogenation catalyzed by Raney-Nickel according to the method reported by De Brabander et al (27) and Jain K et al (28). Briefly, 2.7 ml of ethylenediamine were mixed with 12 ml of acrylonitrile and 26 ml of deionized water. An exothermic reaction occurs, which indicates that double Michael addition reaction has started. The system was placed at 80°C for one hour using a temperature-controlled heating plate. Under these conditions, the double Michael addition reaction is allowed to complete. Excess acrylonitrile was removed as azeotropic water by rotoevaporation at 40°C and 16 mbar for 15 minutes. The final Michael reaction product corresponds to the intermediate generation dendrimer EDA-dendri (CN)_4_ 0.5 G. The product obtained was dissolved in methanol, placed in the catalytic hydrogenation vessel and completed by volume with 2.5 g of Raney-Nickel as catalyst, and deionized water. The final volume of the reaction vessel was 70 ml. The catalytic hydrogenation reaction was carried out at 24 bar pressure, 70°C, 200 rpm and a constant flow of gaseous hydrogen for 1 hour. Excess water was removed by rotoevaporation at 80°C and 16 mbar for 15 minutes. This product corresponds to the dendrimer of PPI 1.0 G. The whole synthesis process was repeated until dendrimers in higher generations were obtained 4.0 G and 5.0 G. For the synthesis of higher generations, the amount of acrylonitrile was increased. The generational growth process of the PPI dendrimers is depicted in Fig 1.

**Fig 1.**
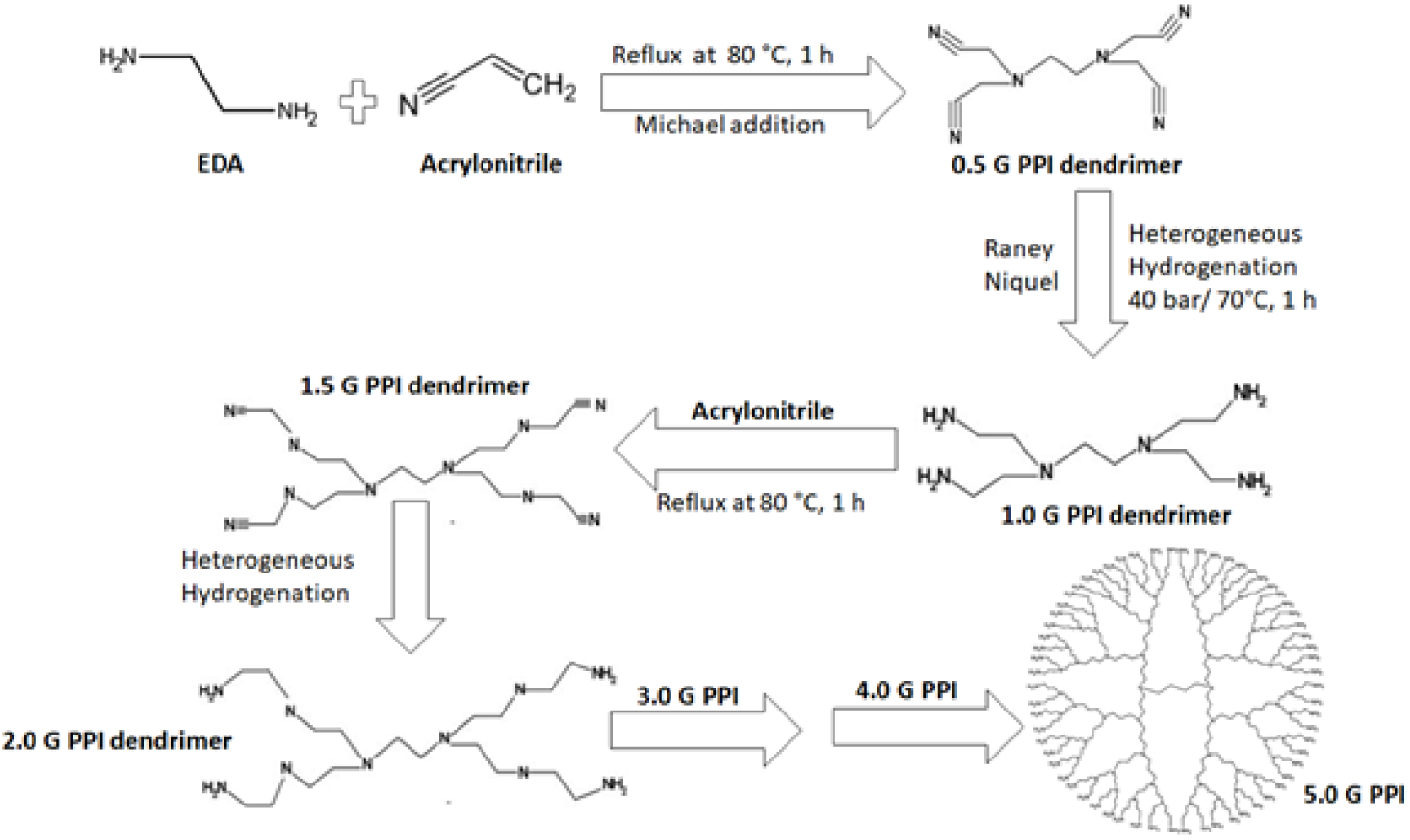
Generational growth process of poly(propylene imine) PPI dendrimers.

### 2.4 Physicochemical and morphological characterization of PPI dendrimers

We determined the topography of the dendrimers using the transmission electron microscopy (TEM), FEI Tecnai G2 F20, including a sample preparation system. Samples were stained with uranyl acetate as a contrast medium. We performed chemical characterization using Fourier Transform Infrared Spectroscopy (FTIR) and proton nuclear magnetic resonance (^1^HNMR). A Thermo Scientific Nicolet iS50 infrared spectrometer, Massachusetts, USA, was used. The samples were placed directly in sample holders of the equipment. The FTIR spectra were recorded at room temperature, in a working range of 4000 - 400 cm^-1^ with a resolution of 4 cm^-1^. The ^1^HNMR technique was performed to confirm the synthesis of PPI dendrimers. The samples were analyzed in a Bruker AMX400 spectrometer, Texas, USA. We dissolved the samples in deuterated chloroform and analyzed them at 300 MHz. The size of the nanoparticles and the concentration of nanoparticles per milliliter of the PPI dendrimers were determined using the Nanoparticle Tracking Analysis (NTA) technique. To perform the analyses, the samples were dispersed in deionized water at a ratio of 1:5. Additionally, we characterized the drug encapsulated in the PPI dendrimer by FTIR and TEM of under the same conditions as mentioned above.

### 2.5 Benznidazole encapsulation in PPI dendrimers

We encapsulated Benznidazole in PPI 5.0 G dendrimer using the equilibrium dialysis method. We weighed and dissolved 50 mg of the dendrimer in 50 ml of ultrapure water. The solution was placed under constant stirring at 200 rpm and 37°C using a CORNING PC-420D magnetic stirring and heating plate. 39 mg of the pure drug was added to the dendrimer dissolved in water. The dendrimer solution with drug was left in constant stirring at 37°C for 72 hours. Finally, it was dialyzed using a standard MWCO cellulose membrane between 12-14 kDa and 29 mm in diameter. The solution was placed in the dialysis bed and immersed in 250 ml ultrapure water. The dialysis process was carried out for 60 minutes at 170 rpm and 37°C. At the end of the dialysis time, the membrane was removed and the product, corresponding to the drug encapsulated in the dendrimer, was collected. We determined the encapsulation efficiency of the trypanocidal drug by indirect method, calculating the amount of non-encapsulated drug, for which an aliquot of 1 ml of dialysis water was taken, filtered through a 0.45 µm membrane, placed in the vial and analyzed by HPLC. Finally, we calculated indirectly the loading efficiency and the percentage of encapsulated drug, using the following mathematical relationships:

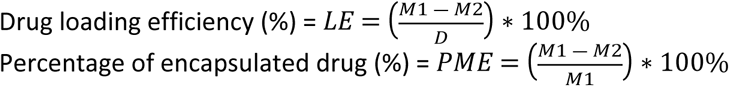

### 2.6 Benznidazole release from PPI 5.0 G dendrimers in vitro

We determined the drug release kinetics in intestinal simulation fluids. For this purpose, we used a

0.01 M Phosphate-Buffered Saline (PBS) solution adjusted to pH 7.4, suitable for cell culture. In vitro release was performed using the equilibrium dialysis technique. We took 5 ml of the encapsulated drug solution from the dendrimer and placed on cellulose membrane (12-14 kDa). The cellulose membrane was sealed at both ends and immersed in 25 ml of release medium. The system was stirred at 400 rpm and 37°C for 240 hours. We withdraw 1 ml of sample and replace it with 1 ml of fresh medium at known time intervals. We filtered the removed aliquot on 0.45 µm pore membrane and analyzed by HPLC. Finally, we calculated the concentration of the drug released from the standardized calibration curve for the pure drug. For comparison purposes, we performed in vitro release of the pure drug. For this purpose, 20 mg of BZN were weighed, placed on the dialysis membrane, and dialyzed in PBS, according to the method described above. We performed the release of pure BZN during 48 hours.

## 3. Results and Discussion

### 3.1 Synthesis and characterization of PPI dendrimers in 4.0 and 5.0 G generation

Synthesis of PPI-type dendrimers was achieved by double Michael addition reaction followed by catalytic hydrogenation. Ethylenediamine was used as the core of the dendrimer and the acrylonitrile molecule contributed the branches to the dendrimer growth. PPI dendrimers grow radially and in generations, starting at the 1.0 G generation and reaching up to the 5.0 G generation. During synthesis, intermediate generations are formed first, which have -CN groups in their structure; these groups are transformed into primary amines - NH_2_ by heterogeneous hydrogenation catalyzed by Raney Niquel (29). During the synthesis, a color variation from pale yellow, in the 1.0 G generation, to yellowish brown, in the 5.0 G generation, was evidenced. The synthesized dendrimers presented viscous consistency, similar to bee honey, except for the intermediate generation 0.5 G, which appeared itself as a white solid mass. The physicochemical and morphological characterization performed by FTIR evidenced the presence of 4.0 G and 5.0 G generation dendrimers. Fig 2 shows the FTIR diffractogram of the PPI 5.0 G dendrimer. In the infrared analysis, a rocking vibration characteristic of the CH_2_ bonds was observed at 796 cm^-1^; at 1063 cm^-1^, the bending vibrations of the N-H bonds of the tertiary amines are observed; at 2936 cm^- 1^, the asymmetric tension vibrations of the C-H bonds are present; at 1670 cm^-1^, a narrow band corresponding to the vibrations of the N-H groups is observed; a band of lower intensity was observed at 1414 cm^-1^, which corresponds to the scissoring vibrations of the CH_2_ groups. A narrow band at 2550 cm^-1^, indicates the self-tension of the nitrile C ≡ N groups and finally around 3436 cm^- 1^ a broad and pronounced band corresponding to the characteristic tension vibration of the NH_2_ bonds of the terminal amino groups is observed. The peaks obtained confirmed that the nitrile groups were reduced to amino terminal groups. Additionally, the FTIR results were confirmed by ^1^HNMR, as shown in Figure 2-b. Between 0.9 and 1.3 ppm, the methyl group (CH_2_) peaks coming from the dendrimer core were obtained and the alkyl groups -CH_2_-CH_2_-were obtained between 1.2 and 1.65 ppm, between 2.548 ppm and 2.693 ppm, the peak of the tertiary amine protons [-N (CH_2_)_3_] is found; between 2.712 and 2.994, the primary amine groups (CH_2_NH_2_) are found and finally a pronounced peak is observed at 7.307, which corresponds to the primary amine proton, thus confirming the synthesis of PPI dendrimer at generation 5.0 G. It is noteworthy that the peak at 7.307 ppm in the PPI 5.0 G dendrimers is more pronounced than in the PPI 4.0 G dendrimers, (data not shown in the paper) due to the higher number of hydrogens in the dendrimer structure. While the dendrimer in generation 4.0 G has 32 amino terminal groups, the dendrimer in generation 5.0 G has 64 amino terminal groups. Similar bands in both FTIR and ^1^HNMR were present in the PPI 4.0 G dendrimer. The results found are similar to those reported by Patel et al. (30) and by Birdhariya et al. (31). Fig 3 shows the morphology of the dendrimers in 4.0 G and 5.0 G generation. The dendrimers are of radial growth, the morphology is spherical in the higher generations, and therefore, in generation 4.0 G a spherical morphology can be appreciated, with some irregularities and trying to form agglomerates. As the number of terminal amino groups increases, the shape is completely spherical and with no tendency to agglomerate, as can be seen in the micrograph of the dendrimer in 5.0 G generation. According to the literature, one of the special properties of dendrimers is their spherical shape and nanometer size, these morphological properties make them excellent candidates for hydrophobic drug nanocarriers. According to Sonam Choudhary et al. (2017), the dendrimers present compact and globular structure with spherical shape and regular architecture (21), a description that agrees with the results obtained in the synthesis of PPI dendrimers in higher generations by divergent approach. Table 1 shows the particle size corresponding to PPI 4.0 G and 5.0 G dendrimers. According to the results obtained for the size of the dendrimers by NTA, it is evident that as the dendrimer grows in generation, its size increases. The obtained size growth results agree with the dendrimer size comparison study performed by Prashant Kesharwani et al. In this study, they compared by TEM and DLS analysis the size of PPI dendrimers in 3.0 G, 4.0 G and 5.0 G generation (32).

**Table 1.** Size and concentration of PPI dendrimers in 4.0 G and 5.0 G generation

| PPI dendrimer generation | Size (nm) |  | Dendrimer concentration/ml |
| --- | --- | --- | --- |
|  | Average | SD |  |
| PPI 4.0 G | 146 | 63 | $5.40 \times 10^8 \pm 7.79 \times 10^7$ |
| PPI 5.0 G | 157 | 66 | $1.02 \times 10^9 \pm 2.78 \times 10^7$ |

**Fig 2.**
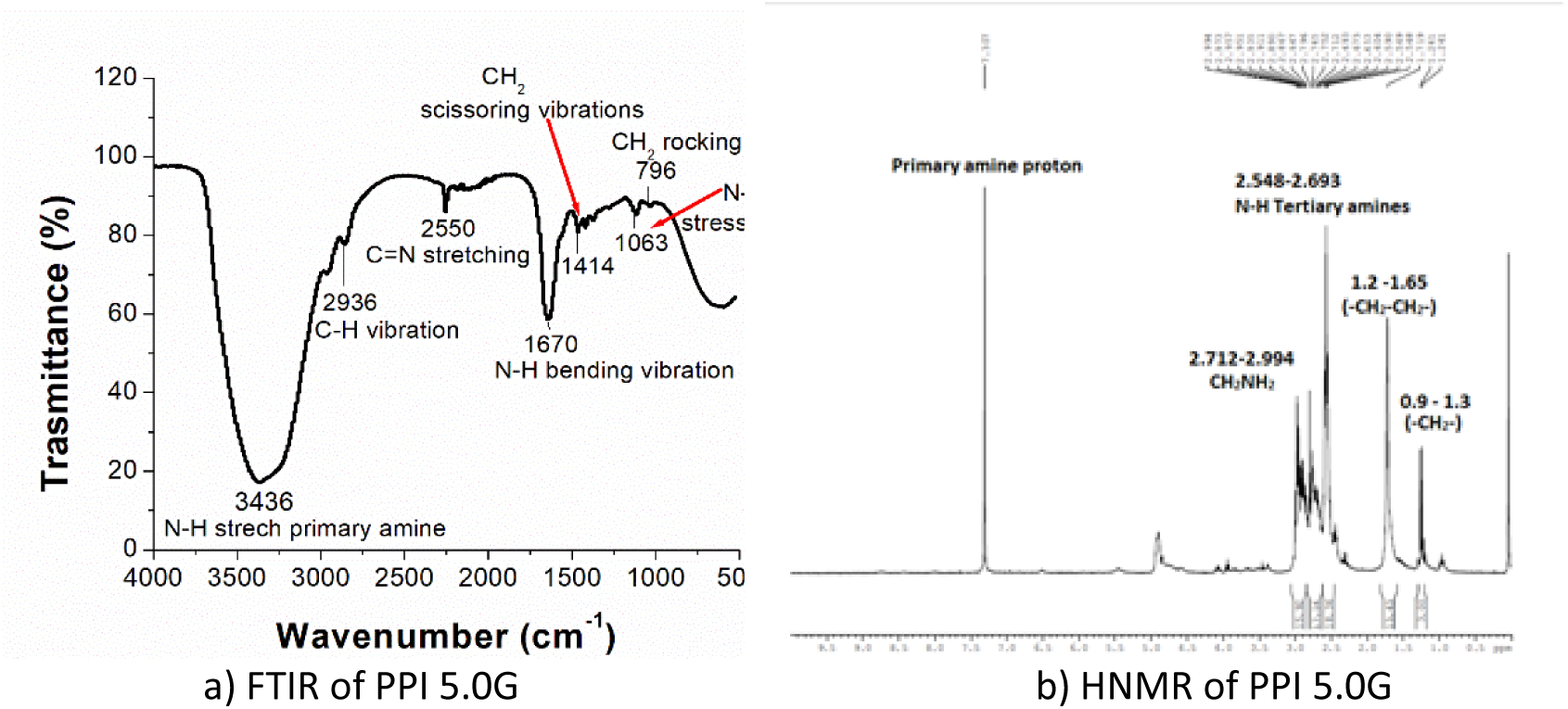
Chemical characterization of PPI dendrimers in 5.0 G generation

**Fig 3.**
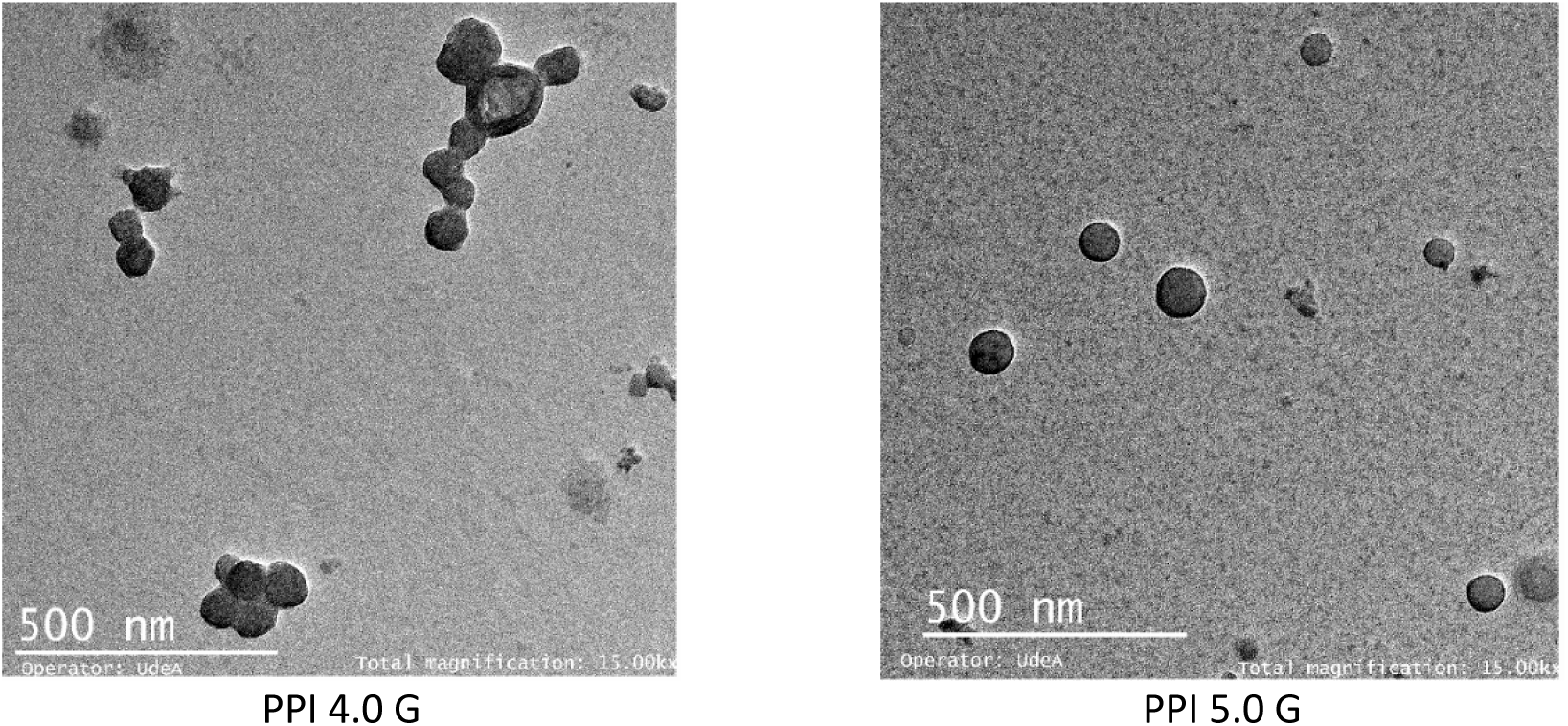
Morphology of PPI dendrimers in 4.0 G and 5.0 G generation synthesized by divergent approach.

### 3.2 Determination of encapsulation efficiency and entrapment efficiency of BZN in PPI 5.0 G dendrimers

The entrapment efficiency and encapsulation efficiency of BZN was performed indirectly. Two drug concentrations, 10 µM and 30 µM, were evaluated. The concentrations were determined from the standardized calibration curve for the pure analyte. The results obtained are shown in Table 2.

**Table 2.** Loading and entrapment efficiency of Benznidazole in PPI 5.0 G dendrimers

| Encapsulated drug | BZN mass (mg) | Loading efficiency (%) | Entrapment efficiency (%) |
| --- | --- | --- | --- |
| BZN/PPI 5.0G 30 μM | 39 | 78 | 99,61 |
| BZN/PPI 5.0G 10 μM | 13 | 17,6 | 67,69 |

As shown in Table 2, increasing the amount of BZN to be encapsulated increases the encapsulation efficiency and the drug entrapment efficiency at this loading percentage, which is close to 100% of drug molecules. This indicates that the hydrophobic BZN molecules adhere to the dendrimer cores via hydrophobic and hydrogen bonding interactions. The dendrimer cores are formed of tertiary amines; therefore, they tend to retain the BZN particles, through hydrogen bridge type interactions. In addition, the cavities of the PPI dendrimer are highly hydrophobic, which increases the possibility of interaction with hydrophobic drugs such as BZN (33). According to the literature, the amount of host molecules entrapped in the dendrimer is proportional to the shape and size of the molecule to be encapsulated, as well as to the shape and size of the available internal cavities of the dendritic structure (34). In addition, higher generations of dendrimers have greater capacity and in turn more space to encapsulate hydrophobic fractions (22), allowing for improved solubility properties of drugs such as BZN, which has low solubility and low permeability (35). Considering the above, the synthesized PPI 5.0 G dendrimers are suitable structures for encapsulating BZN. These structures presented adequate internal cavities to house the drug molecules, evidenced by the entrapment percentage of 99.61% and the loading efficiency of 78%. Entrapment efficiency is an important parameter in determining the drug release characteristics from the dendrimer (36). The drug was physically entrapped within the dendritic structure due to the presence of spherical cavities. These cavities are hydrophobic and exhibit affinity for drugs with similar solubility characteristics, such as BZN. Likewise, the drug can form hydrogen bridges with the nitrogen atoms present in the PPI cavities (37). Therefore, the interactions of PPI towards BZN are non-covalent, hydrophobic and Van der Walls type interactions, mainly hydrogen bonds. Non-covalent type interactions are used to improve the solubility of insoluble drugs (38).

In addition, drug encapsulation was confirmed by FTIR and NTA techniques. Figure 4 shows the FT-IR spectrum of the drug encapsulated in the PPI 5.0 G dendrimer.

**Fig 4.**
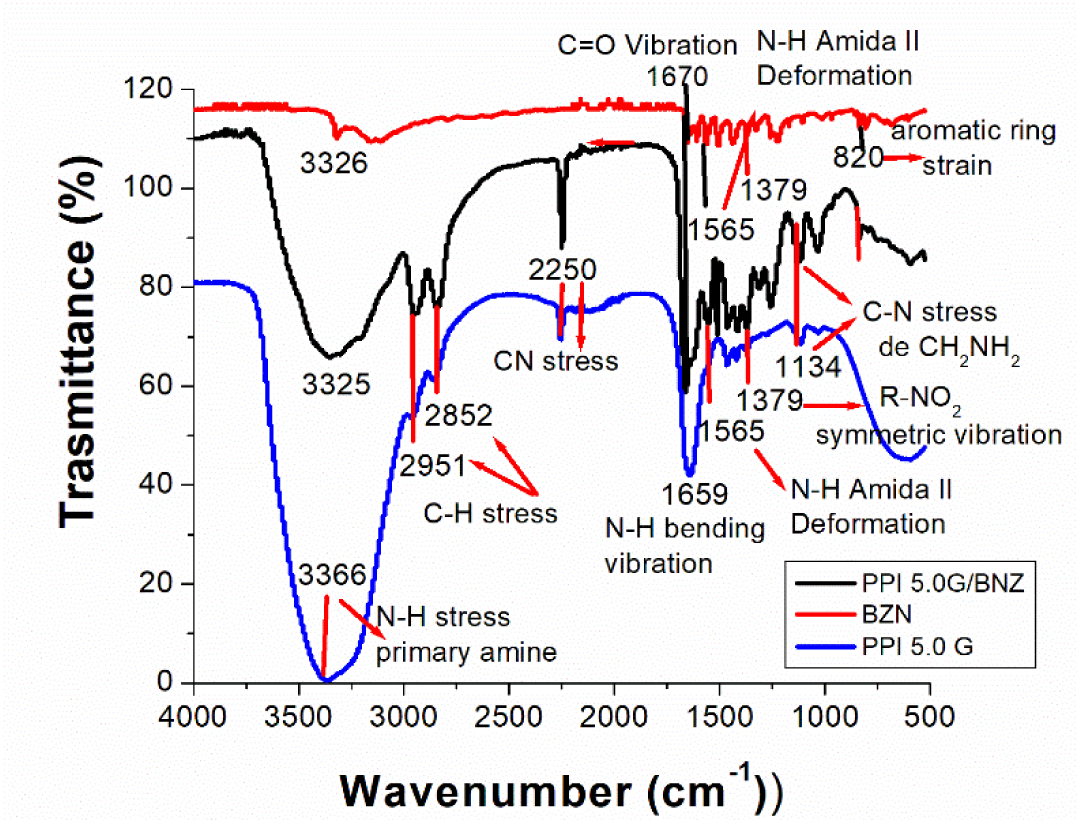
Comparison of the FT-IR spectra of PPI 5.0 G (black line), BZN without PPI (red line), and BZN encapsulated in PPI 5.0 G (blue line) The FT-IR spectrum of the encapsulated drug shows the characteristic bands of BZN and PPI 5.0 G. This demonstrates that the drug was successfully encapsulated in the dendrimer. The IR spectrum of the drug can be divided into three fragments: imidazole group, benzyl group, and the acetamide fragment. It shows an intense band close to 3281cm^-1^ characteristic of the absorption of the secondary amines present in the acetamide fragment (39). This band overlaps with the characteristic bands of the primary amines of the PPI dendrimer, which occur in the region of 3550 cm^-1^ and 3320 cm^-1^. The amide I band was observed at 1565 cm^-1^, while the amide II bands showed N-H bending strains at 1565 cm^-1^. The vibration at 1670 cm^-1^ is characteristic for the bond of the carbonyl group C=O of the drug. The tension bands in the aromatic ring appear at 820 cm^-1^. The intensity of the band in the tension of the C-N bond present in the imidazole group was observed at 1157 cm^-1^. The R-NO_2_ functional group showed characteristic symmetric vibration at 1379 cm^-1^. The values found in the characteristic bands are in agreement with those reported in the literature (39). The NTA assay confirmed an increase in the size of the dendrimer with the encapsulated drug. The free dendrimer had a size of (157 ± 66) nm, while the dendrimer containing the encapsulated drug had a size of (194± 47) nm.

### 3.3 In vitro release kinetics of encapsulated Benznidazole in PPI 5.0 G dendrimers

The estimation of the release profile was performed in vitro, by the equilibrium dialysis technique, using 0.01M PBS and pH 7.4 as the release medium. Fig 5 and fig 6 show the release behavior of BZN over time. Fig 5 indicates the release profile in mg/ml concentration and Fig 6 shows the release percentage. From the figures, it can be inferred that there was a controlled and sustained release over time. The maximum amount of drug released was at 230 hours. It is important to note that there were no abrupt releases of the drug; on the contrary, the behavior of the curve was of slow growth over time.

**Fig 5.**
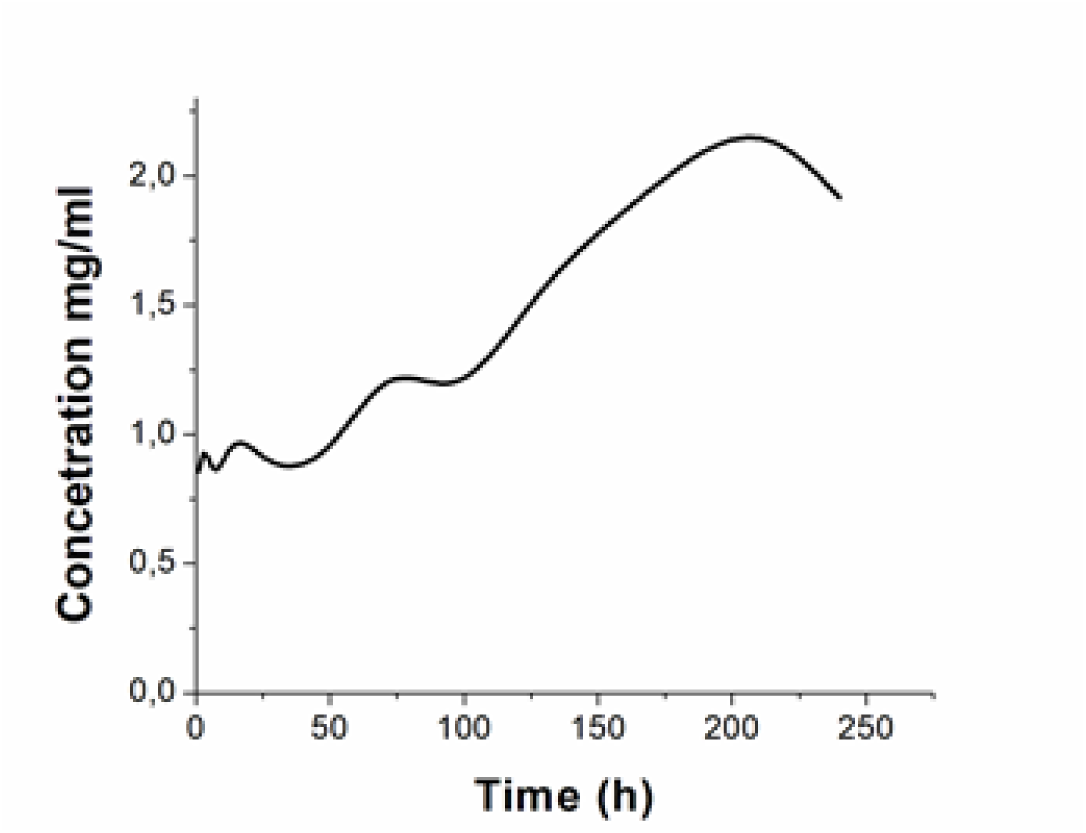
BZN release from PPI 5.0G dendrimer at concentration.

**Fig 6.**
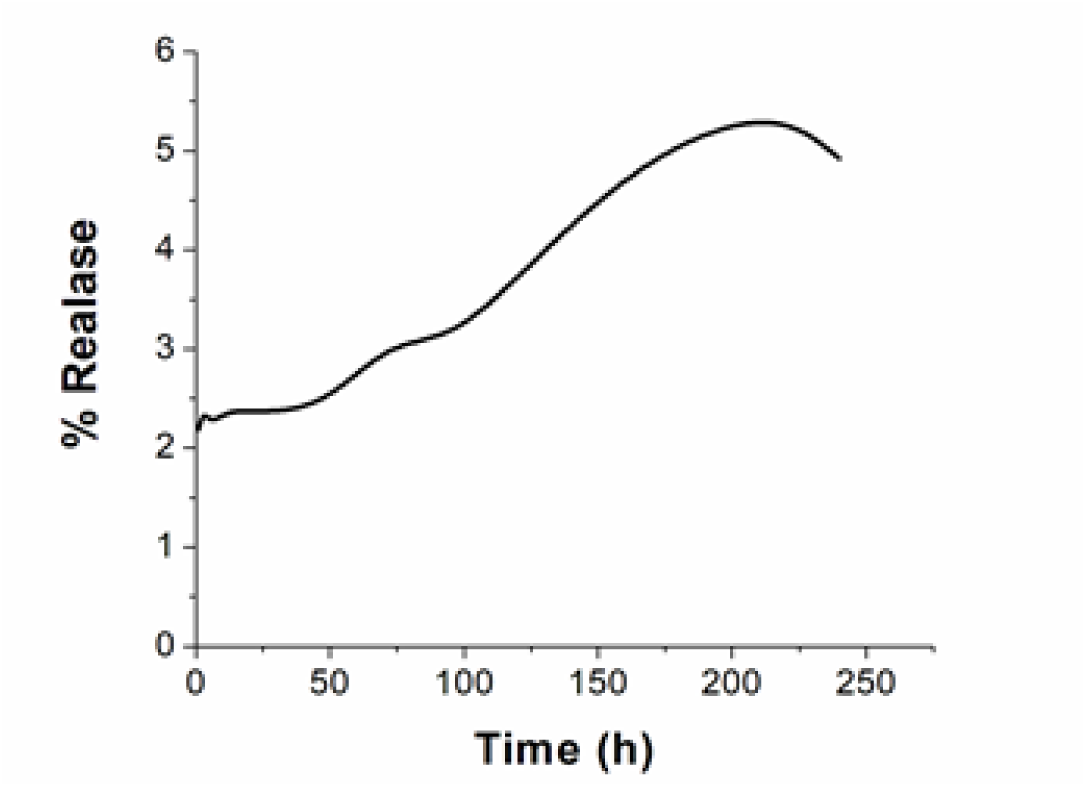
BZN release curve of PPI 5.0 G dendrimer.

Fig 6 shows the release of dendrimer-free BZN in 0.01M PBS and pH 7.4. The release was carried out for 48 hours. From the figure, we can see that release is fast. In the first 24 hours, about 40% of the drug was released. Compared to fig 5, it was observed that the entrapped drug in the dendrimer does not allow the BZN molecules to be easily released. This release behavior is related to the entrapment efficiency of the nanostructure towards the drug, which was 99.61%. Fig 6 shows that 2.5% of the drug was released in the first 48 hours, compared to Fig 7, which shows that 40% of the drug was released in the same hours. Evidencing that dendrimers as drug transporters improve the pharmacokinetic properties of drugs, improve their solubility in aqueous media, remain longer in the blood circulation, improve transit through biological barriers and delay drug degradation (24).

**Fig 7.**
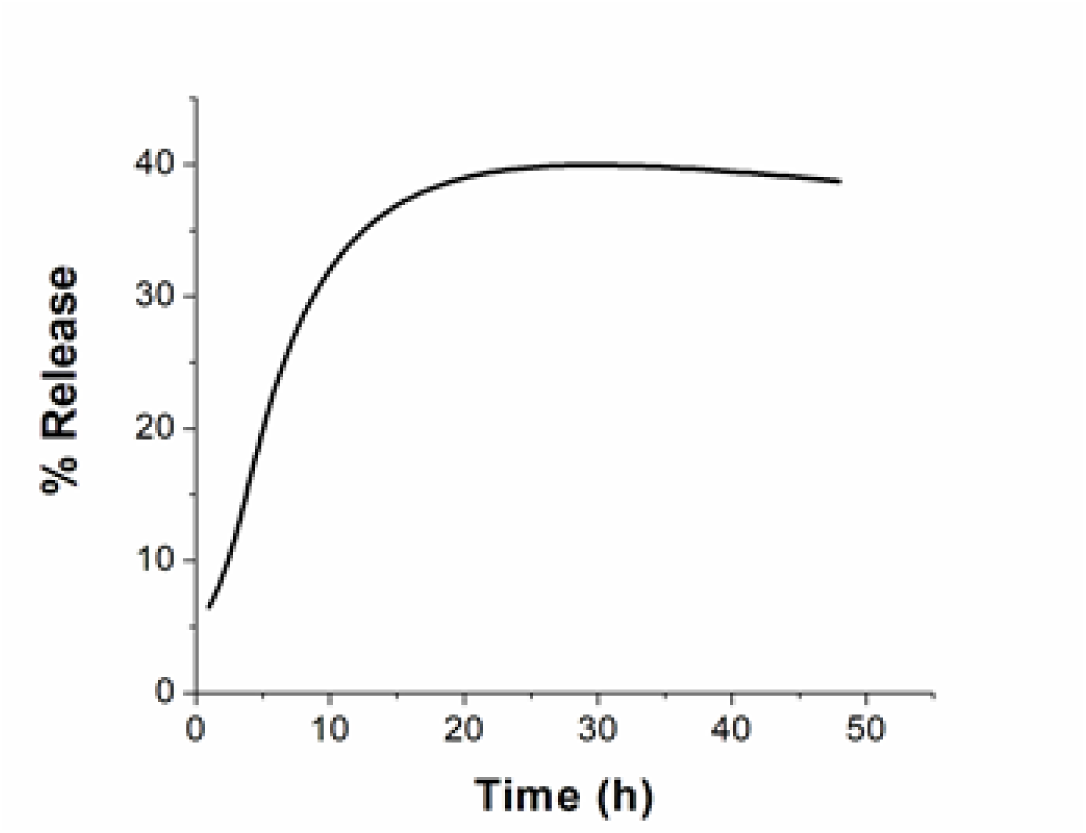
Release curve of BZN in PBS, pH 7.4.

## Conclusion

In the present research, it was possible to synthesize PPI dendrimers in 4.0 G and 5.0 G generation. The physicochemical and morphological characterization confirmed the synthesis of these nanostructures. The synthesized dendrimers presented spherical shape and nanometric size, favorable characteristics to encapsulate hydrophobic drugs. The synthesized PPI dendrimers were used to encapsulate BZN. The drug was encapsulated with a loading efficiency of 78% and an entrapment efficiency of 99.6%. The release kinetics of the PPI 5.0 G - BZN system showed a prolonged and sustained release profile over time. Using PPI 5.0 G dendrimers to encapsulate drugs such as BZN could be an alternative for the treatment of Chagas disease since it would improve the physicochemical properties of the drug.

## Acknowledgements

The authors thank Dr. Isabel Cristina Ortiz Trujillo, from the Biología de Sistemas group at UPB, for allowing us to use the laboratories and facilities to perform the experiments. Also, to Dr. Luis Alberto Ríos from the Industrial Chemical Process Group of the Universidad de Antioquia, where the materials used in this work were synthesized.

## Funding

The author Jenny Ordoñez-Benavides was supported by Ministerio de Ciencia Tecnología e Innovación – Minciencias (call 647), Ph.D. studies. This work was supported by the UPB-Innova Program at Universidad Ponticia Bolivariana. The publication fees were financed by the Centro de Investigaciones para el Desarrollo Integral - CIDI of the Universidad Ponticia Bolivariana UPB, and the Research Operations Department of the Instituto Tecnológico Metropolitano de Medellín ITM.

## Notes

### Competing Interest Statement

The authors have declared no competing interest.

